# Improving the Z_3_EV promoter system to create the strongest yeast promoter

**DOI:** 10.1101/2024.05.24.595832

**Authors:** Rina Higuchi, Yuri Fujita, Hisao Moriya

**Author notes:** Competing interest The authors declare that there is no competing interest in this work. Author contributions RH performed experiments, RH and YF analyzed the data, and RH and HM wrote the paper.

## Abstract

Promoters for artificial control of gene expression are central tools in genetic engineering. In the budding yeast *S. cerevisiae*, a variety of constitutive and controllable promoters with different strengths have been constructed using endogenous gene promoters, synthetic transcription factors and their binding sequences, and artificial sequences. However, there have been few attempts to construct the highest-strength promoter in yeast cells. In this study, by incrementally increasing the binding sequences of the synthetic transcription factor Z_3_EV, we were able to construct a promoter (P36) with approximately 1.4 times the strength of the *TDH3* promoter. This is stronger than any previously reported promoter. Although the P36 promoter exhibits some leakage in the absence of induction, the expression induction by β-estradiol is maintained. When combined with a multicopy plasmid, it can express up to approximately 50% of total protein as a heterologous protein. This promoter system can be used to gain knowledge about the cell physiology resulting from the ultimate overexpression of excess proteins and is expected to be a useful tool for heterologous protein expression in yeast.

## Introduction

Promoters play a central role in the artificial control of gene expression (Carey and Smale 2000). In the budding yeast *S. cerevisiae*, various promoters have been constructed using endogenous gene promoters (Romanos, Scorer, and Clare 1992; Weinhandl et al. 2014; Rajkumar et al. 2016; Peng et al. 2015), synthetic sequences (Vaishnav et al. 2022), synthetic transcription factors and their binding sequences (Azizoglu, Brent, and Rudolf 2021; McIsaac et al. 2011; Gligorovski, Sadeghi, and Rahi 2023). These promoters are characterized and utilized based on differences in expression strength, whether they are constitutive or controllable, and other factors. For controllable promoters, the method of control (such as temperature, drugs, light, etc.), the controllability (signal-to-noise ratio, minimum and maximum strength, etc.), and convenience (whether endogenous inducers are present or whether the promoter and control factor need to be introduced simultaneously) are important considerations. As for endogenous promoters, the *TDH3* promoter, known for its maximum expression strength, and the *GAL1* promoter, which can be repressed by glucose and induced by galactose, have been commonly used (Peng et al. 2015). Recently, there has been active development of synthetic promoters using artificial transcription activators/repressors and their binding sites. Examples include the WTC_846_ system (Azizoglu, Brent, and Rudolf 2021), which incorporates tet0 sites into the *TDH3* promoter and integrates feedback control of transcription factors, achieving tetracycline inducibility, high signal-to-noise ratio, wide dynamic range, and strong maximum expression strength; a promoter system that uses multiple binding sites for the synthetic transcription factor Z_3_EV (McIsaac et al. 2011, 2013, 2014), which can be induced by β-estradiol and avoids gratuitous transcription induction; and light-inducible promoter systems (Gligorovski, Sadeghi, and Rahi 2023). These promoters have been continuously improved to meet various criteria required for gene expression control. However, efforts to maximize expression strength have been limited.

In this study, we aimed to construct a promoter specifically designed to maximize the expression of recombinant proteins in yeast. Previously, we used the *TDH3* promoter to maximize the expression of excess proteins (Eguchi et al. 2018; Namba et al. 2022). However, the *TDH3* promoter had issues with its still insufficient strength, concerns about transcriptional competition with endogenous promoters especially when used on multicopy plasmids, and decreased expression in the post-diauxic phase due to its role as glycolytic proteins. To overcome these issues, we focused on the following conditions: 1) the promoter itself must have the highest transcriptional activation activity; 2) when the promoter is incorporated into a multicopy plasmid, the expression of other endogenous genes should not be affected by competition with transcription factors; 3) unintended proteins should not be expressed due to gratuitous induction; 4) the promoter should be minimally affected by the growth phase and do not need to change from the optimal growth conditions (i.e. glucose can be used as a primary carbon source). We thus considered that the Z_3_EV system promoter would match these conditions (McIsaac et al. 2013, 2014). The Z_3_EV system promoter is an artificial promoter constructed by modifying the *GAL1* promoter. This promoter is regulated by the synthetic transcription factor Z_3_EV and is induced by the drug β-estradiol. Z_3_EV is a composite protein consisting of a zinc-finger DNA-binding domain, the estrogen (β-estradiol) receptor, and the VP16 transcriptional activation domain. When β-estradiol binds to Z_3_EV, it moves into the nucleus and induces expression as a synthetic transcription factor (McIsaac et al. 2011). Because it is a completely synthetic system, it is expected that there will be a minimal reduction in the expression of other genes due to competition with transcription factors, and almost no gratuitous protein expression (McIsaac et al. 2013). Particularly, when integrating the comparative studies of the strongest promoters conducted so far (McIsaac et al. 2014; Kotopka and Smolke 2020; Gligorovski, Sadeghi, and Rahi 2023), the P3 promoter with six binding sites for the Z_3_EV promoter was considered the strongest. Attempts to increase the strength of this promoter using random mutagenesis with machine learning were made, but they were not very successful (Kotopka and Smolke 2020). On the other hand, attempts to increase the number of Z_3_EV binding sites have not been made. Therefore, in this study, we attempted to increase the strength by incrementally increasing the number of binding sites. As a result, the incremental increase in the number of Z_3_EV binding sites (up to 12) led to an increase in expression strength, achieving approximately 1.43 and 1.25 times the strength of the *TDH3* promoter and the P3 promoter, respectively. Increasing the number of binding sites beyond this point reduced the expression level. Although this promoter exhibited increased leakage in the absence of induction, transcriptional activation by β-estradiol was still maintained.

Using this promoter with the combination of the gTOW multicopy plasmid system (Moriya et al. 2012; Moriya, Shimizu-Yoshida, and Kitano 2006), we were able to express up to approximately 50% of the total protein as a heterologous protein. This promoter system can thus be used to gain knowledge about the cell physiology resulting from the ultimate expression of excess proteins and is a useful tool for heterologous protein expression in yeast.

## Results

### The P3 promoter exhibits strength comparable to the *TDH3* promoter

Because the P3 promoter has been reported to have strength equivalent to that of the *TDH3* promoter (Kotopka and Smolke 2020) we first verified whether the strength of the P3 promoter is indeed equivalent to that of the *TDH3* promoter using a fluorescent protein reporter assay. Specifically, we introduced the low-toxicity green fluorescent protein moxGFP (Namba et al. 2022) downstream of each promoter and constructed the pTOW plasmid (**Figure 1A, B**). The pTOW plasmid utilizes the low-expression leucine synthesis enzyme gene (*leu2-89*) as a marker which works as a selection bias to increase the plasmid copy number >100, allowing for maximum protein expression of the target protein on the plasmid when cultured in leucine-deficient SC–LeuUra medium (Moriya et al. 2012; Moriya, Shimizu-Yoshida, and Kitano 2006). Transcription from the P3 promoter is induced by 1 μM β-estradiol. **Figure 1C** shows the time-course changes in fluorescence and growth of each strain measured with a fluorescence plate reader. The maximum growth rate and maximum fluorescence level of each strain are shown in **Figures 1D and 1E**. As expected, the *TDH3* promoter exhibited constitutive GFP expression with or without β-estradiol, while the P3 promoter showed significant GFP expression induction upon β-estradiol addition. Moreover, the maximum expression under the SC–LeuUra conditions was comparable to that of the *TDH3* promoter (**Figure 1E)**. Additionally, under SC–Ura conditions, the expression from the P3 promoter was significantly higher than that from the *TDH3* promoter (**Figure 1D)**. Therefore, it was confirmed that the P3 promoter has strength equal to or greater than that of the *TDH3* promoter.

**Figure 1.**
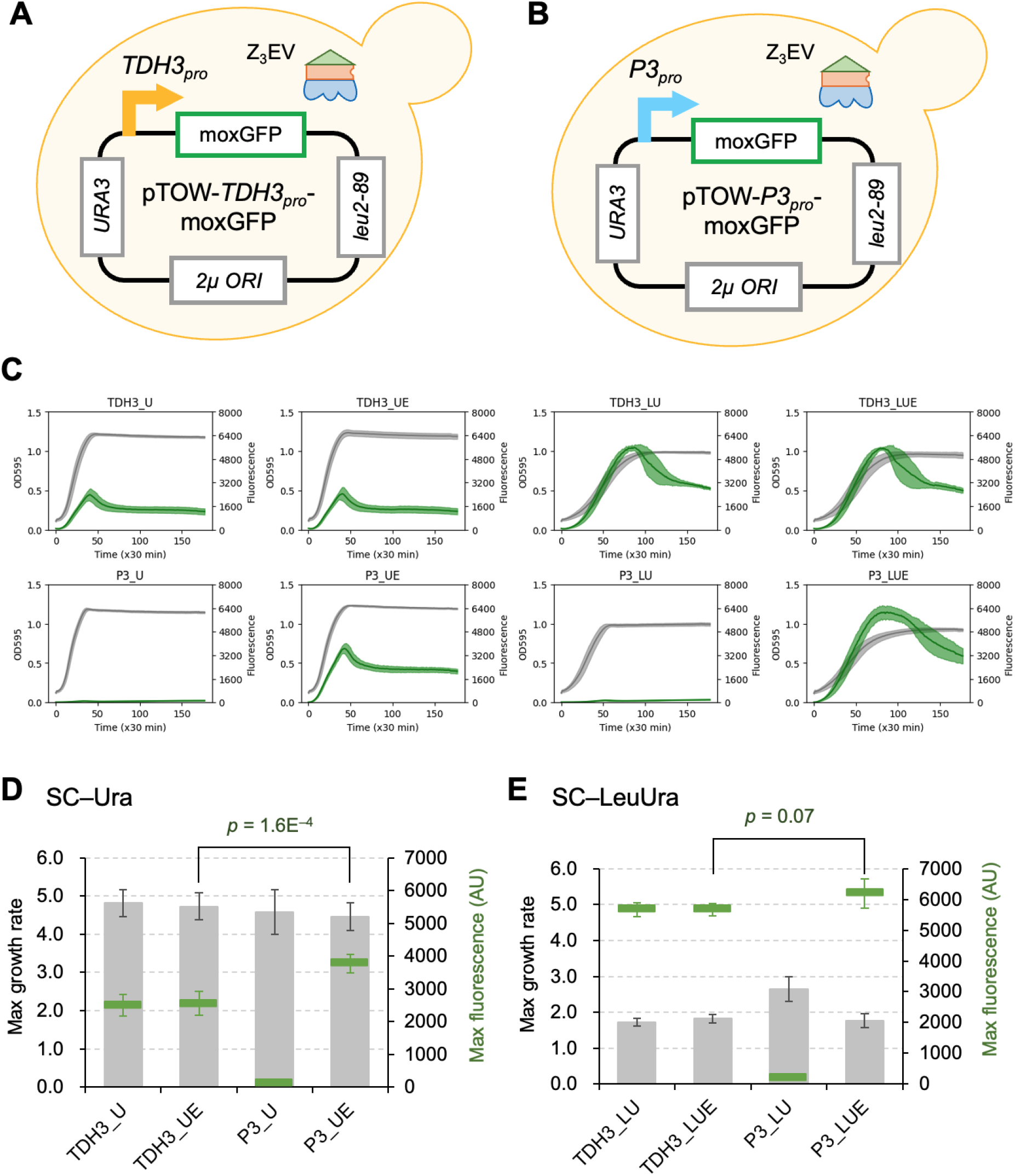
The P3 promoter exhibits strength comparable to the *TDH3* promoter. **A, B**) Schematic diagrams of the cells used in this study. We used strains of DBY12394, which express the transcription factor Z_3_EV necessary for the P3 promoter, transformed with pTOW plasmids. MoxGFP expression is driven by the *TDH3* promoter (*TDH3*_*pro*_, **A**), or the P3 promoter (*P3*_*pro*_, **B**). **C**) Growth curves and fluorescence values over time. Gray indicates the growth curve, and green indicates the fluorescence curve. The shaded areas represent the standard deviation. The left axis shows the turbidity of the culture measured at OD595, and the right axis shows the fluorescence intensity of moxGFP. **D, E**) The maximum growth rate and max fluorescence intensity calculated from **C**. The gray bar graph represents the max growth rate, and the green marker graph represents the max fluorescence intensity, with each error bar indicating standard deviation. Statistical tests were performed using Welch’s t-test (two-tailed). Measurements in **C-E** were performed using a fluorescence plate reader. In **C, D**, and **E**, “U”, “LU”, “UE”, and “LUE” represent SC–Ura, SC–LeuUra, SC–Ura+β-estradiol, and SC–LeuUra+β-estradiol, respectively. β-estradiol was added at a concentration of 1 μM. Experiments were performed with four or more biological replicas.

### A promoter with 12 Z_3_EV binding sites (P36) shows the highest transcriptional activity

Z_3_EV-based promoters, including the P3 promoter, can change their strength by altering the position and number of Z3EV binding sites on the promoter (McIsaac et al. 2014). Therefore, we attempted to construct a promoter stronger than the *TDH3* promoter by increasing the number of Z_3_EV binding sites in the P3 promoter. Using the P3 promoter with 6 Z_3_EV binding sites as the base, we constructed new promoters with 4 binding sites, which is 2 fewer (P3-2), and incrementally added 2 binding sites at a time, up to a maximum of 16 binding sites (P3+n, where n is 2 to 10) (**Figure 2A**). The constructed promoters were inserted into the pTOW plasmid and their strength was evaluated using a reporter assay, as described above. The results are shown in **Figure 2B-E**.

**Figure 2:**
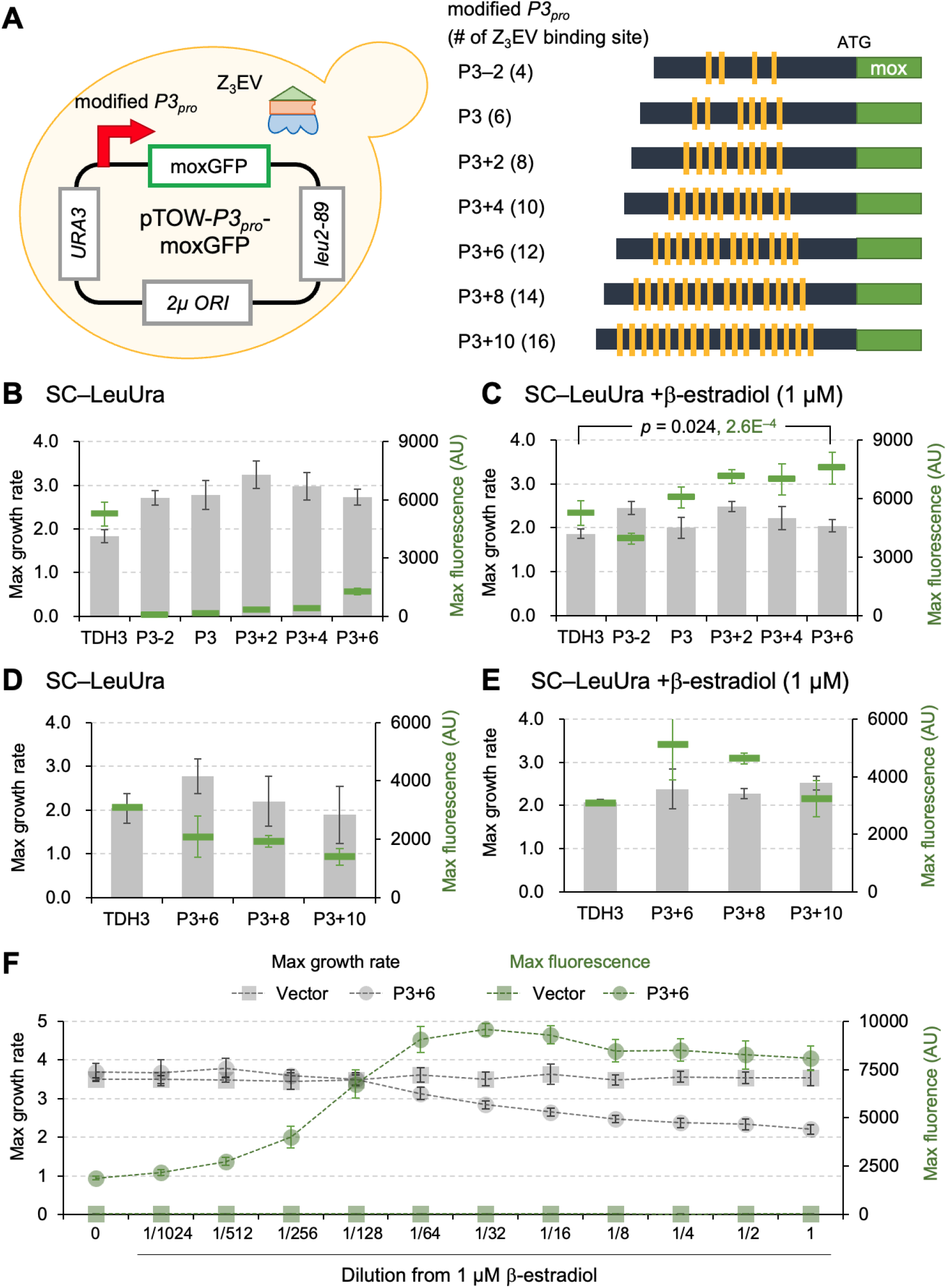
The effect of increasing Z_3_EV binding sites on transcriptional strength. **A**) A schematic diagram of the cell used in the experiment and the structures of the modified P3 promoters. Orange blocks indicate the positions of the Z_3_EV binding sequences (gcgtgggcg). **B**-**E**) Maximum growth rate (gray bar graph) and maximum fluorescence intensity (green markers) in strains harboring plasmids with each promoter in SC–LeuUra medium (**B** and **D**) and SC–LeuUra medium with 1 μM β-estradiol added (**C** and **E**). In **C**, black text indicates the *p*-value for the maximum growth rate, and green text indicates the *p*-value for the maximum fluorescence intensity. **F**) Induction of expression by β-estradiol for the P3+6 promoter. Maximum growth rate (gray markers) and maximum fluorescence intensity (green markers) in SC–LeuUra medium with β-estradiol (diluted from 1 μM by half) are shown. **B**-**E** experiments show the mean values and standard deviations (error bars) of four biological replicates measured by a fluorescence plate reader. Statistical tests were performed using Welch’s t-test (two-tailed).

As shown in **Figure 2C**, under the maximum expression condition (SC–LeuUra+β-estradiol), the maximum fluorescence intensity increased with the increase in the number of Z_3_EV binding sites, and the P3+6 promoter was confirmed to have 1.43 and 1.25 times the strength of the *TDH3* promoter and the P3 promoter, respectively. Additionally, as shown in **Figure 2B**, even under non-induced conditions (SC–LeuUra), the maximum fluorescence intensity increased with the increase in the number of Z_3_EV binding sites, indicating leakage expression with the increase in binding sites. On the other hand, when the number of Z_3_EV binding sites was increased to 14 (P3+8) and 16 (P3+10), the maximum fluorescence intensity decreased (**Figure 2E**). From these results, it was suggested that the P3+6 promoter with 12 Z_3_EV binding sites is the strongest among the promoters constructed by modifying the P3 promoter. We note that the maximum expression level from the P3+6 promoter was higher than that of the *TDH3* promoter (*p* = 2.6E^-4^), while the maximum growth rate at this time was higher with the P3+6 promoter than with the *TDH3* promoter (*p* = 0.024, **Figure 2C**). Therefore, it is suggested that the P3+6 promoter imposes a less extraneous burden, such as transcription factor competition, which the *TDH3* promoter potentially carries.

Next, we investigated the inducibility of the P3+6 promoter by β-estradiol (**Figure 2F**). As predicted from **Figure 2B**, there was leakage expression even in the absence of β-estradiol, but the expression level increased with the rise in β-estradiol concentration from zero ((Max fluorescent = 1893) to 1/32 (Max fluorescent = 9562). Therefore, the inducibility was maintained with a maximum induction/non-induction ratio of 5.1. The addition of β-estradiol at concentrations of 1/64 (16 nM) or higher caused a decrease in cell growth. The fact that such growth reduction was not observed in the vector control, and that the degree of growth reduction became stronger with increasing β-estradiol concentration, suggests that the expression level increased stepwise from concentrations above 1/64 to 1 μM, causing growth inhibition due to the associated burden. Therefore, by using the P3+6 promoter (hereafter referred to as the P36 promoter), it is possible to investigate the effects on cells due to stepwise increases in expression levels, particularly in high-expression regions.

### Maximum expression of excess protein using the P36 promoter

In our recent study, we found that mox-YG, a mutated form of moxGFP that loses fluorescence, can be expressed in the highest amounts within yeast cells **(Fujita et al**., ***in preparation*)**. The promoter used in that study was the *TDH3* promoter. We then tested whether even more protein could be expressed using the P36 promoter. In **Figure 3A**, the results of SDS-PAGE analysis of total proteins in cells during the logarithmic growth phase (OD660 = 0.9-1.1) in SC–LeuUra medium with the addition of 1 μM β-estradiol are shown. In **Figures 3B** and **3C**, the quantitative results of the expressed proteins are shown as the ratio of the target protein amount to the total protein amount in the vector control. The moxGFP expression level from the P36 promoter is 1.4 times that of the *TDH3* promoter, which closely matches the measurement of promoter strength by fluorescence (**Figure 2C**). As expected, mutations that cause the loss of fluorescence increased protein expression levels in both promoters. Furthermore, expression from the P36 promoter was higher than from the *TDH3* promoter, achieving up to approximately 50% mox-YG expression. The total protein amount at this time was not different from the vector control (**Figure 3C**, p = 0.70, Welch’s t-test (two-tailed)). Therefore, the increase in mox-YG expression is directly reflected in the decrease in the amount of other proteins within the cell.

**Figure 3:**
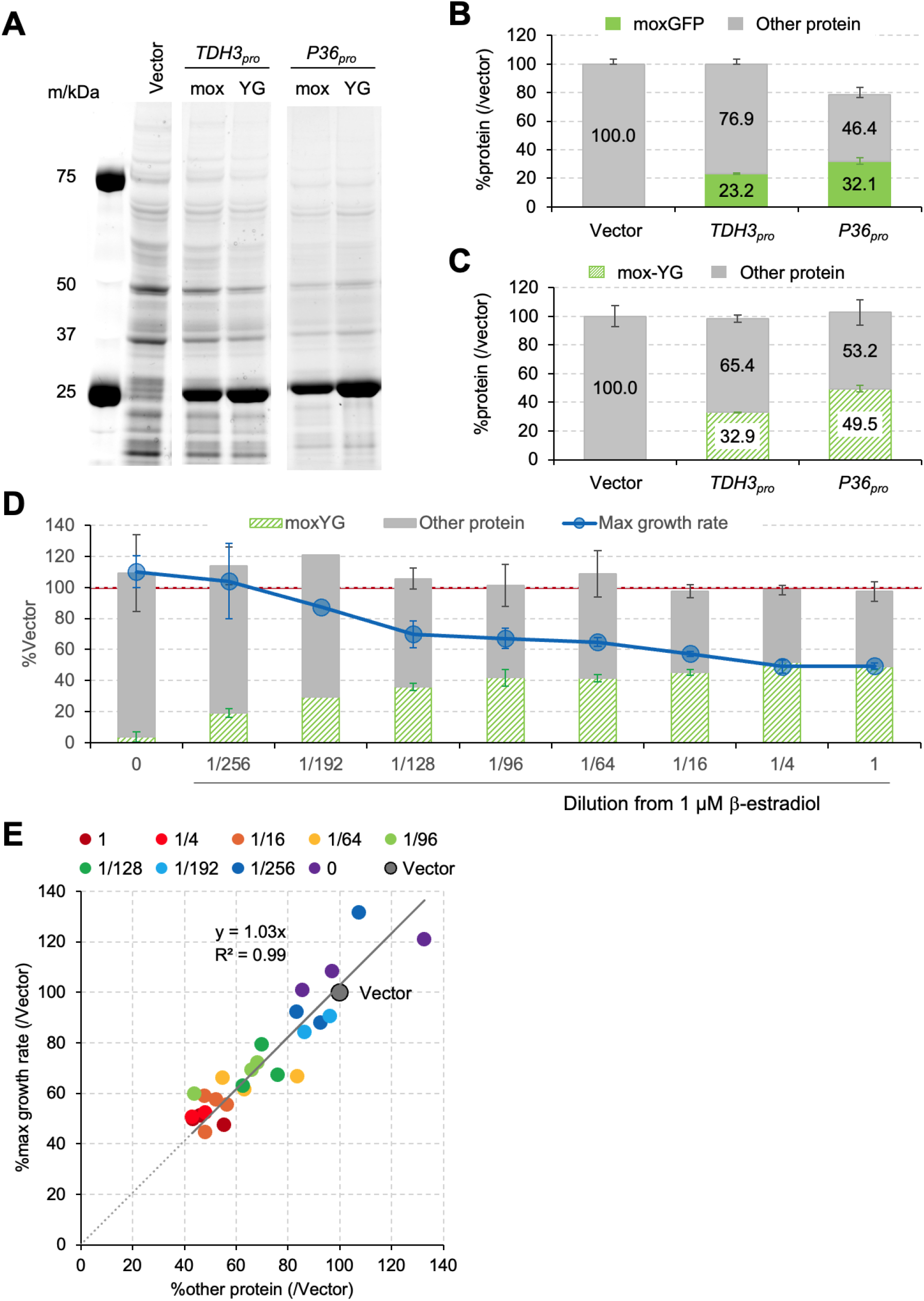
Maximum expression of excess protein using the P36 promoter. **(A**) Image of total proteins separated by SDS-PAGE. mox: moxGFP, YG: mox-YG. (**B, C**) Protein amounts measured from the SDS-PAGE gel images. The expression levels of moxGFP, mox-YG, and other proteins (Other protein) were calculated based on the total protein amount of the vector control, which was set to 100. The bar graphs show the average values obtained from three biological replicate experiments, with error bars indicating the standard deviation. (**D**) Maximum growth rate, mox-YG, and other protein amounts in cells expressing mox-YG from the P36 promoter with a stepwise dilution of β-estradiol. β-estradiol was added to SC-LeuUra medium in dilutions starting from 1 μM. The graph shows the averages of three biological replicates. The mean values and standard deviations (error bars) were calculated from three biological replicates, except for the 1/192 dilution, whose mean was calculated from two biological replicates. The red line indicates the total protein amount (100%) in vector control cells. (**E**) The relationship between maximum growth rate and protein amount in cells expressing mox-YG from the P36 promoter with a stepwise dilution of β-estradiol. All three biological replicates are shown as individual points, except for the 1/192 dilution (shown with two biological replicates). The regression line, its regression equation, and the R^2^ value on a graph when performing linear regression through the origin are shown. For **A**-**E**, the cultures and OD660 measurements were performed using a small shaking culture device

From previous experiments with *E. coli* and yeast, it is known that the increase in expression of excess proteins causes a gradual decrease in growth (Scott et al. 2010; Kafri et al. 2016). Therefore, we measured the growth rate and protein amount when the induction of mox-YG was gradually strengthened (**Figure 3D, E**). As a result, strengthening the induction of mox-YG gradually decreased the growth rate. During this time, the total protein amount, including both mox-YG and other proteins, remained almost unchanged. Therefore, as the expression of mox-YG increased, the amount of other proteins decreased. Under these conditions, the decrease in maximum growth rate and the decrease in the amount of proteins other than mox-YG (Other protein) could be approximated as a straight line through the origin with a slope close to 1 (1.03), and the R^2^ value was 0.99 (**Figure 3E**). From these results, it was found that the P36 promoter has the maximum strength to reach 50% of the total protein and allows for gradual regulation of protein expression.

## Discussion

In this study, we aimed to construct an artificial promoter with high expression strength by increasing the number of Z_3_EV binding sites. The P36 promoter, which has six additional binding sites in the P3 promoter, was the strongest among those constructed, with a strength 1.4 times that of the *TDH3* promoter and 1.2 times that of the P3 promoter (**Figure 2C**). Although there was significant leakage without β-estradiol addition, the inducibility was maintained at approximately fivefold (**Figure 2F**). By combining this promoter with the multi-copy gTOW system, it was possible to massively express the harmless protein mox-YG to about 50% of the total protein. When the number of binding sites was further increased, a decrease in strength was observed (**Figure 2D, E**). The reason for this decrease could be that the Z_3_EV binding sites are too close to each other or to the transcription start site, causing interference. Different configurations of the binding sites might further increase promoter strength.

The P36 promoter is considered to be a very powerful tool for studying the growth inhibition effects (protein burden) caused by the overproduction of excess proteins. In *E. coli* studies, there is a linear relationship between the expression of excess proteins and growth reduction, with an estimate that expressing around 37% excess protein results in growth cessation (Bruggeman et al. 2020). A similar linear relationship has been observed in yeast cells, but the amount of excess protein that can be expressed was not as high (Kafri et al. 2016). One of the reasons for this is considered to be the insufficient strength of the promoters. Using the P36 promoter developed in this study, it is possible to express up to 50% excess protein (**Figure 3C**), and through stepwise induction, a wide range of linear relationships similar to those seen in *E. coli* were observed (**Figure 3E**). Interestingly, yeast cells expressing 50% excess protein maintained a 50% growth rate, with the origin of the regression line being zero (**Figure 3E**). This suggests that eukaryotic cells have a higher capacity to accommodate excess proteins compared to prokaryotic cells and that prokaryotic and eukaryotic cells may have different response regimes to protein burden.

## Materials and Methods

The reagents, strains, plasmids, and primer sequences used in this study are summarized in the Key Resource Table.

### Yeast growth conditions and transformation

The budding yeast strain DBY12394 (*MATα ura3Δ leu2Δ0::ACT1pro-Z*_*3*_*EV-NatMX*) was used (McIsaac et al. 2013). Yeast transformation was performed using the lithium acetate method (Amberg et al. 2005). Cells were cultured in synthetic complete (SC) medium (Amberg et al. 2005) without uracil (–Ura) or leucine and uracil (–LeuUra). All cultures were maintained at a temperature of 30 °C.

### Plasmids Used in this study

Plasmids used in this study are listed in the Key Resource Table. In constructing the plasmid, synthetic DNA and PCR-amplified DNA were joined using the recombination-based method in yeast (Oldenburg 1997), and their structures were verified by DNA sequencing. All plasmids are high-copy plasmids with a *2μ ORI* and carry the low-expression *LEU2* allele (*leu2-89*). Therefore, after introducing these plasmids into *LEU2*-deficient strains and culturing in leucine-depleted media (SC–LeuUra), the plasmid copy number increases to over 100 copies. Due to the principle of the genetic tug-of-war (gTOW), the copy number of the plasmids increases to the level at which the target protein causes growth inhibition (Moriya et al. 2012; Moriya, Shimizu-Yoshida, and Kitano 2006).

### Measurement of Growth and Fluorescence

The promoter expression strength was evaluated using a reporter assay with moxGFP as the fluorescent protein reporter. Yeast cells were cultured statically under their respective medium conditions. For the measurements, a microplate reader (TECAN Infinite F200) was used to monitor and measure OD595 and Ex 485 nm/Em 535 nm every 30 minutes. The maximum growth rate was determined as described in the previous study (Moriya, Shimizu-Yoshida, and Kitano 2006).

### Protein Analysis and Quantification

Yeast cells overexpressing the target protein were pre-cultured in SC–LeuUra medium, and then cultured in 5 mL of SC–LeuUra medium with or without β-estradiol using a shaking culture apparatus (ADVATEC, TVS062CA). Cells in the logarithmic growth phase (OD660 = 0.9-1.1) were treated with 1 mL of 0.2 N NaOH (Kushnirov 2000), followed by total protein extraction using 100 μL of LDS sample buffer (ThermoFisher). For each analysis, total protein was extracted from the amount of cells equivalent to 1.0 OD at OD660 (1 ODu). The extracted total proteins from 0.1 ODu cells were labeled with Ezlabel Fluoroneo (ATTO) according to the manufacturer’s protocol and separated by 4-12% sodium dodecyl sulfate-polyacrylamide gel electrophoresis (SDS-PAGE). Protein detection and quantification were performed using the SYBR-green fluorescence detection mode of the LAS-4000 image analyzer (GE Healthcare) and Image Quant TL software (GE Healthcare). The total protein amount of the vector was set to 100%, and the amounts of moxGFP, mox-YG, and other proteins were quantified.

## Data Availability Statement

Strains and plasmids are available upon request. The authors affirm that all data necessary for confirming the conclusions of the article are present within the article and figures.

## Acknowledgments

We would like to express our gratitude to the members of the Moriya Laboratory at Okayama University. This research was partly supported by the Nagase Science and Technology Foundation, the Grant-in-Aid for Scientific Research (KAKENHI) grant numbers 24K02013, 22K19294, and 20H03242.

